# No One-Size-Fits-All: An Evidence-Based Framework to Select Plasma EV Isolation Methods

**DOI:** 10.64898/2026.03.09.710675

**Authors:** Scheila Julia Werle, Marie Louise Nautrup Therkelsen, Chen Meng, Mads Grønborg, Lise Lotte Gluud, Dres Damgaard

**Author notes:** These authors contributed equally to this work and share first authorship (Scheila Julia Werle, Marie Louise Nautrup Therkelsen). Corresponding author: Dres Damgaard, Novo Nordisk A/S, Novo Nordisk Park 1, 2760 Måløv, Denmark. Data availability statement: Raw LC-MS/MS data and processed search files is available in ProteomeXchange Consortium (http://proteomecentral.proteomexchange.org) on the PRIDE16 partner repository (identifier: PXD075093). MSD and nanoFCM data are available as supplementary data. The python code with the data analysis is available in the GitHub repository at https://github.com/mlnth/EV_isolation_methods, released under MIT licence. Funding statement: This research was funded by Novo Nordisk A/S.

## Abstract

Extracellular vesicles (EVs) hold significant promise as biomarkers, but their clinical translation is constrained by variability in pre-analytical handling and isolation. EV isolation methods directly shape which EV populations are captured and characterized, yet systematic method comparisons across multiple analytical dimensions are limited. We comprehensively evaluated eleven EV isolation methods to define their performance and applications. EVs were quantified by NanoFCM, profiled for tetraspanins (CD9, CD63, CD81) via MSD assays, and further characterized by LC-MS/MS proteomics. We show that different EV isolation methods recover different EV populations. Our data provide guidance on method selection based on downstream application needs and serve as a look-up tool if a protein of interest is detected.

EV isolation methods broadened proteome coverage but showed divergent performance and recover different EV populations. While all methods captured EVs in the 50-150nm range, centrifugation and ultracentrifugation identified the broadest proteomes (up to 1093 proteins) driven by higher plasma protein carryover. Conversely, ExoEasy and qEV 70 isolated larger EVs and achieved stronger depletion of abundant plasma proteins but showed lower proteome coverage. A total of 117 proteins were detected across all isolation methods. Pre-clearing samples removed contaminants but at the cost of protein identifications. We demonstrate that method selection must align with the specific analytical goal: centrifugation for comprehensive proteome profiling, affinity/size-exclusion methods for contaminant-sensitive assays, and precipitation for high-throughput applications. This systematic characterization provides an evidence-based framework and look-up resource for matching isolation strategies to downstream applications and research questions.

**Graphical Abstract for Table of Contents:** 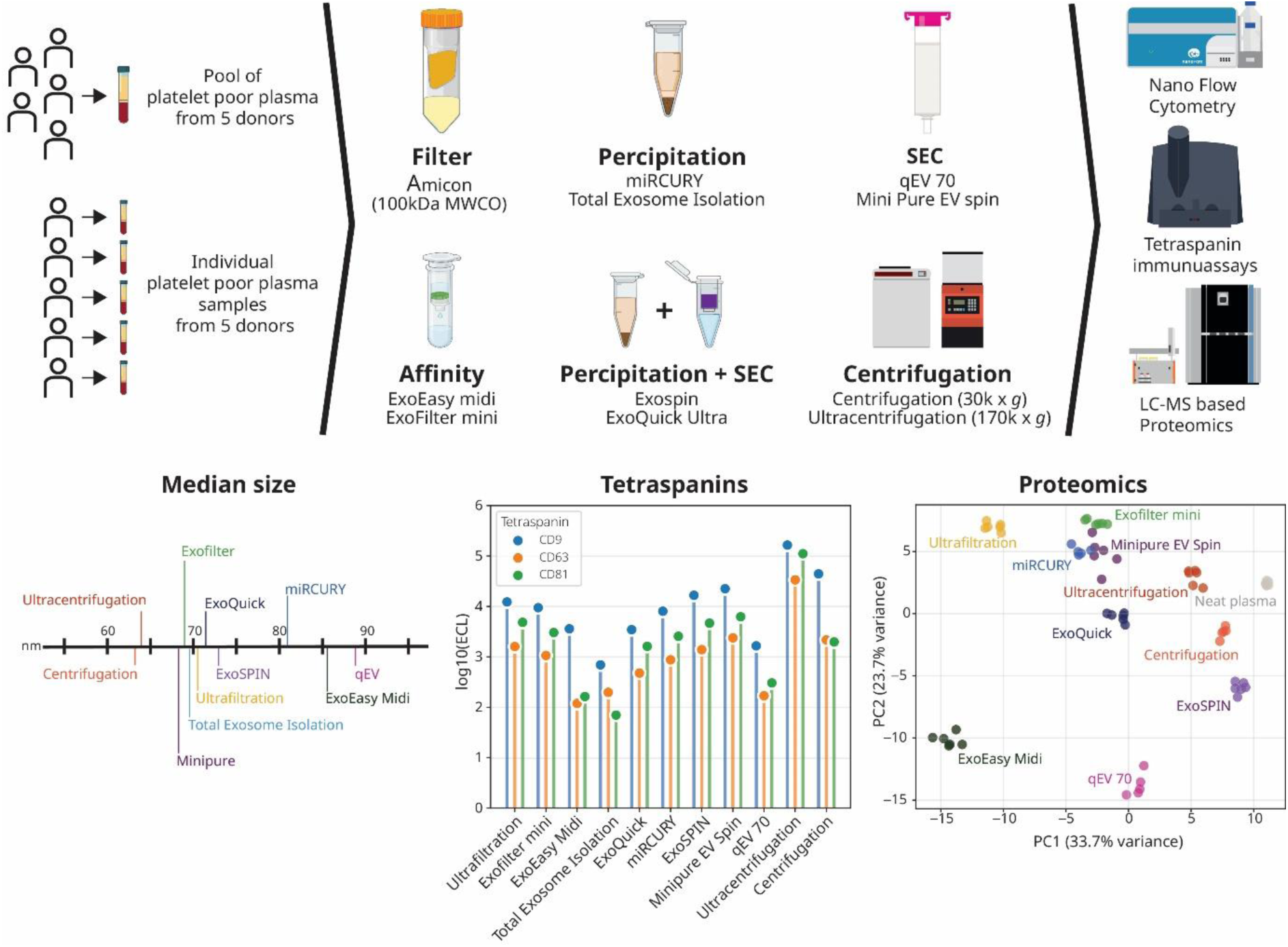

This study evaluated 11 extracellular vesicle (EV) isolation methods which enriched distinct EV subpopulations with varying degrees of contaminants. No single approach optimized purity or proteome coverage; in this paper we present an Evidence-Based Framework to select plasma EV isolation methods based on downstream application needs.

## Introduction

Extracellular vesicles (EVs) are membrane-enclosed particles secreted by cells, serving as molecular fingerprints of their cellular origin and indicators of pathophysiological changes. These nano-sized particles can be broadly classified into exosomes, microvesicles, and apoptotic bodies, each with distinct biogenesis and characteristics. Their stable presence in biological fluids combined with diverse molecular cargo that may include proteins, nucleic acids, and metabolites, makes them promising for biomarker purposes. Additionally, their availability in accessible fluids such as blood, saliva, and urine enables minimally invasive disease monitoring. Their diagnostic and prognostic potential spans across multiple conditions, including cancer, liver diseases, cardiovascular, neurodegenerative, and inflammatory conditions [1, 2, 3, 4, 5]. Inconsistent isolation protocols and analytical variations across laboratories lead to inconsistent findings, hampering clinical application. Standardized, transferable methodologies are urgently needed and the successful implementation of EVs for clinical evaluations depends on robust isolation and characterization methods. [6, 7, 8].

To address these challenges, we focused on manufacturer-recommended protocols with minimal modifications, aiming to establish reproducible and transferable methods. While recent work has evaluated EV isolation methods using minimal plasma volumes [9], most clinical settings readily accommodate larger blood draws, allowing full utilization of manufacturer-recommended input volumes to maximize method efficiency and EV recovery. We therefore conducted a systematic comparison of eleven isolation methods based on different principles including centrifugation, ultrafiltration, membrane affinity, size-exclusion chromatography, and precipitation using blood derived samples. We further investigated the impact of pre-analytical variables, including blood collection as well as plasma pre-centrifugation and pre-filtration, selectively modulate the EV proteome by influencing the enrichment and depletion of specific protein subsets. These methods vary in their scalability, equipment needs and processing time– factors critical for clinical applicability and large-scale implementation.

We characterized the isolated EVs through multiple analytical approaches: Nano Flow cytometry for particle size and concentration [10, 11], MSD immunoassay for tetraspanin evaluation, and Liquid chromatography-tandem mass spectrometry (LC-MS/MS) proteomics to evaluate method-specific proteome profiles [12, 13, 14]. We evaluated yield, purity, reproducibility and proteome depth to identify optimal methods for specific downstream applications, particularly proteomics, while considering practical implementation factors. We established a workflow for appropriate method selection to guide future research in this field. This work underscores that the optimal isolation method will vary significantly depending on the specific analytical purpose ultimately advancing the potential of EVs as a liquid biopsy tool [15, 16, 17, 18]

## Materials and Methods

### Blood collection and processing

Samples were obtained from healthy donors (n=10) through the ‘*Giv Blood collaboration*’, under Danish Ethical Approval (VEK no. H-D-2007-0055). Blood was drawn into BD vacutainer sampling tubes, including K2 EDTA, citrate, and serum coagulate, and centrifuged at 1,811 x *g* for 10 minutes at room temperature (RT) within 45 minutes from sampling. Samples were further centrifuged at 3,000 x *g* for 10 minutes to obtain platelet-poor plasma (PPP) and platelet-poor serum, which was frozen at -20°C.

Prior to EV isolation all samples were filtered through a 0.45 µm filter and centrifuged at 12,000 × *g* for 30 minutes to deplete large vesicles, cell debris, and other contaminants. For pooled samples, PPP from five healthy donors was pooled. The pre-cleared PPP was aliquoted and stored at -20°C.

### Extracellular Vesicles Isolation methods

Pre-cleared PPP was thawed at RT and processed promptly. Replicates were handled concurrently to assess variability. We tested 11 EV isolation methods (Table 1). All commercial methods followed manufacturer protocols unless specified (full protocols are provided in Supplementary – Extended methods). Isolated EV samples were immediately stored at -20°C for further characterization.

**Table 1:** Overview of EV isolation methods evaluated: Methods are grouped by mechanism and commercial methods are listed with vendor and catalogue number

| Type | Method | Vendor | Catalogue # |
| --- | --- | --- | --- |
| **Filtration based methods** | Ultrafiltration (100 kDa MWCO) | Merck | UFC510096 |
| **Affinity based methods** | Exofilter mini | HansaBiomed | 60001 |
|  | ExoEasy Midi | Qiagen | 77144 |
| **Precipitation based methods** | Total Exosome Isolation | Thermo Fisher Scientific | 4484450 |
|  | ExoQuick Ultra | System Biosciences | EQUltra- 20A-1 |
|  | miRCURY | Qiagen | 76603 |
|  | ExoSPIN | Cell Guidance Systems | EX02 |
| **Size Exclusion Chromatography based methods** | Minipure EV spin | HansaBioMed | HBM-mPEVS |
|  | qEV 1 70nm | Izon | ICO-70 |
| **Centrifugation** | Ultracentrifugation (170,000 x *g*) |  |  |
|  | Centrifugation (30,000 x *g*) |  |  |

### NanoFCM

Following instrument calibration as per the manufacturer’s instructions, isolated EV samples were diluted in Ultrapure TE (Thermo Fisher, J75893.AE) for analysis using the Flow NanoAnalyzer (NanoFcm Inc.) in its Exosome setting. The NanoFCM Profession V2.0 software was utilized to assess both particle concentration and size distribution.

### Immunoassays

SECTOR ELISA plates (Mesoscale Discovery, L45SA) were coated with capture antibodies from antibody sets targeting CD9 (F215M-3), CD63 (F215L), CD81 (F215N) (all Mesoscale Discovery), and CD41a (R&D Systems, MAB530A) in Diluent 100 (Mesoscale Discovery, R50AA-4). After 1h incubation at RT and washing (PBS +0.05% Tween20), (Mesoscale Discovery, R52AA-1). Detection utilized corresponding detection antibodies from the same antibody sets in a CD9/CD63/CD81 cocktail diluted in Diluent 53 (Mesoscale Discovery, R52AB-1) for 1h, followed by immediate reading in MSD Gold Read Buffer B (Mesoscale Discovery, R60AM-4) on a Mesoscale Discovery SECTOR Imager (Model HTS).

For disruption experiments, paired samples of isolated EVs were either maintained intact or treated with 0,25% SDS for 1h at RT to disrupt the membrane bilayer. Both samples were analyzed for tetraspanins CD9, CD63, and CD81 using MSD immunoassay.

### LC-MS/MS proteomics

Sample amount was normalized to the yield obtained from 200 µL of plasma input. Samples underwent a washing process using centrifugal filter tubes, Amicon Ultra, 100 kDa MWCO (Merck, Cat# UFC5100) to facilitate buffer exchange to (PBS). A buffer comprising 10% n-Dodecyl β-D-Maltoside (DDM), Trypsin/Lys-C (Promega, Cat# V5071), and 50 mM triethylammonium bicarbonate (TEAB) were added to the filter tube for the lysis of retained vesicles and subsequent protein digestion. Following digestion, peptides were eluted from the filter tubes and acidified before loaded onto Evotips (EvoSep Biosystems, Cat# EV2013) in accordance with the manufacturer’s protocol. LC-MS/MS analysis was performed using an Orbitrap Astral MS coupled with an Evosep One system (EvoSep Biosystems). The samples were analyzed by single-shot narrow-window data-independent acquisition (nDIA) with a 21-minute gradient with an EvoSep 8 cm column (EvoSep Biosystems, Cat# EV1109). Raw LC-MS files were processed with Spectronaut v18 (Biognosys). DIA-MS/MS spectra were analyzed using the spectral library-free direct-DIA algorithm against a human UniProt FASTA database (version 20250203, 20,433 entries), with separate searches conducted for each isolation method.

### Evaluating Preanalytical variations

EVs was isolated using centrifugation (30,000 x *g,* 1h). To test blood collection and processing methods, EVs was isolated from human EDTA plasma, EDTA PPP, citrate plasma, citrate PPP, serum and platelet poor serum. EV surface markers (CD9, CD63, CD81) and platelet marker CD41a were then quantified using MSD immunoassays.

To evaluate the impact of pre-clearing and post-isolation buffer exchange, we tested four pre-clearing conditions: (i) no pre-clearing, (ii) filtration only (0.45 µm), (iii) centrifugation only (12,000 × *g*, 30 min), and (iv) combined filtration and centrifugation. Following EV isolation, each sample was split, with one aliquot subjected to the buffer-exchange using Amicon Ultra filters (100 kDa) and the other processed without washing. All samples were then analyzed by LC–MS/MS.

## Results

### Influence of pre-analytical sample preparation on EV composition

Distinct EV marker profiles were observed across platelet-rich and platelet-poor plasma (EDTA, citrate) or serum (Figure 1A). Platelet-poor EDTA and citrate plasma showed reduced CD41a indicating effective platelet depletion, while retaining EV tetraspanin signals (CD9, CD63). In contrast, serum showed no reduction in platelet or tetraspanin markers. Pre-clearing (0.45 µm filtration and/or centrifugation at 12,000 × *g* for 30 minutes) prior to EV isolation significantly reduced LC–MS/MS protein identifications (Figure 1B). The proteins from pre-cleared samples largely overlapped with those from non-cleared PPP (Figure 1C), indicating that pre-clearing lowers total identifications without introducing unique proteins. While pre-clearing lowered the intensities of platelet and apoptotic body associated proteins (Suppl. Figure 1B), it also lowered the intensities of EV-associated proteins (Suppl. Figure 1A), suggesting a potential loss of EVs. Post-isolation buffer exchange with Amicon filters (100 kDa MWCO) had little effect on the number of identified proteins in samples from pre-cleared conditions. In contrast, for samples from non-cleared PPP, it reduced the number of identifications (Figure 1B)

**Figure 1:**
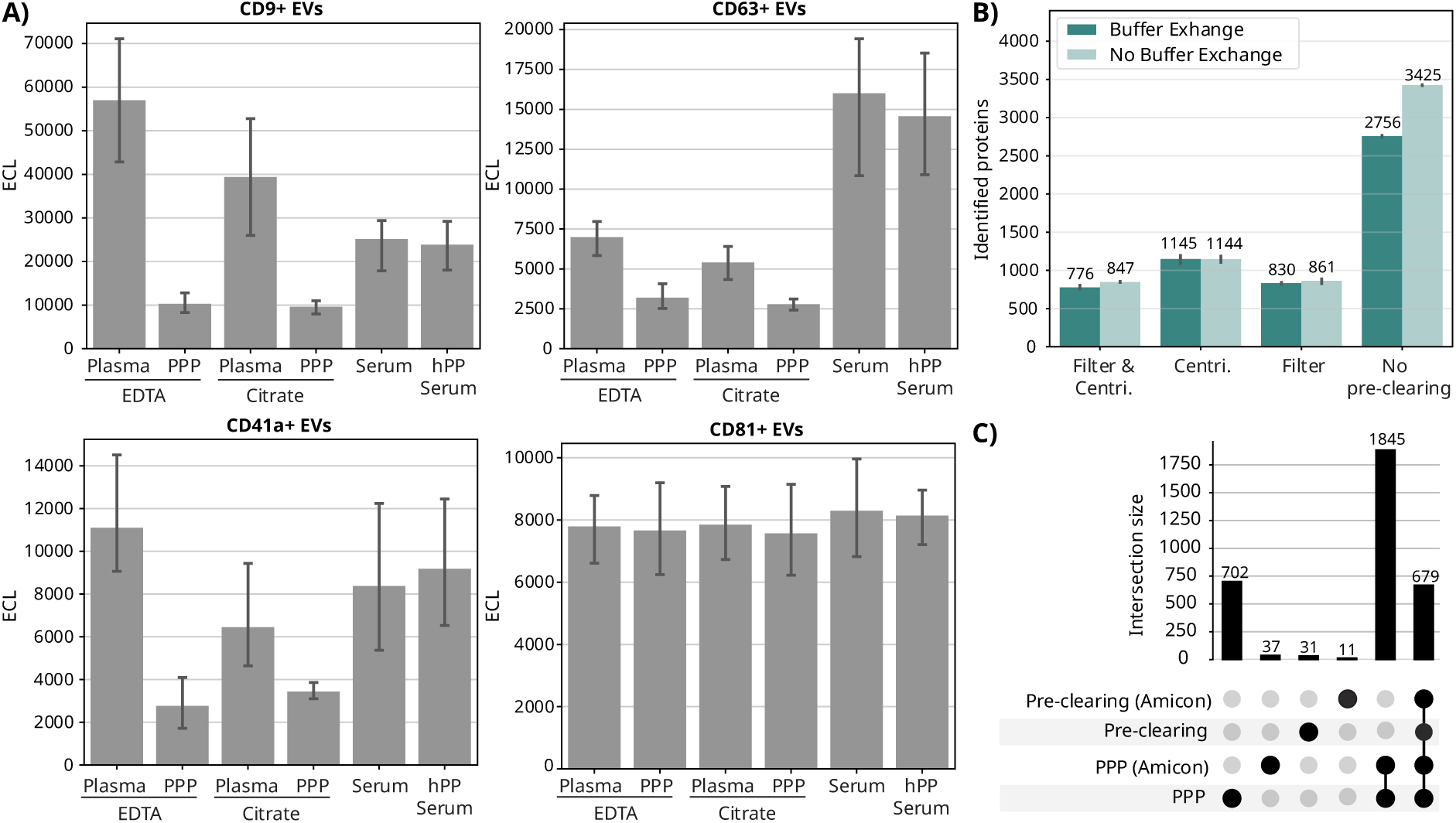
Analysis of preanalytical variations: Analysis on preanalytical variations on EVs isolated with centrifugation (30,000 x *g* for 1 hour). A) Tetraspanin (CD9, CD63, CD81) and platelet marker (CD41a) across different blood collection and processing methods (n=5 donors). Equal amounts of EVs were analyzed by immunoassays. Data points represent individual measurements (ECL). B) Number of proteins identified by LC-MS/MS following EV enrichment with different pre-clearing steps: Centri: centrifugation (12,000 x *g* for 30 min), filtration (0.45 µm), both steps combined, or no pre-clearing. C) Upset plot showing the overlap of identified proteins from pre-cleared PPP or non-pre-cleared PPP, with and without post-isolation buffer exchange with Amicon Ultra filters (100kDA). ECL= Electrochemiluminescence, PPP = platelet poor plasma, h=Human.

### EV characteristics

Principal Component Analysis (PCA) of EV isolation methods (Figure 2A) highlights distinct clustering based on tetraspanin signals, particle yield, and size. Ultracentrifugation stands apart, driven by high tetraspanin levels (Figure 2D-F). Methods like miCURY, ExoQuick, ExoSPIN, Exofilter, Centrifugation, and Minipure group together, mainly influenced by particle yield (Figure 2B). Notably, Total Exosome Isolation and ExoEasy separated by showing high signals after vesicle disruption (Figure 2G-I).

**Figure 2:**
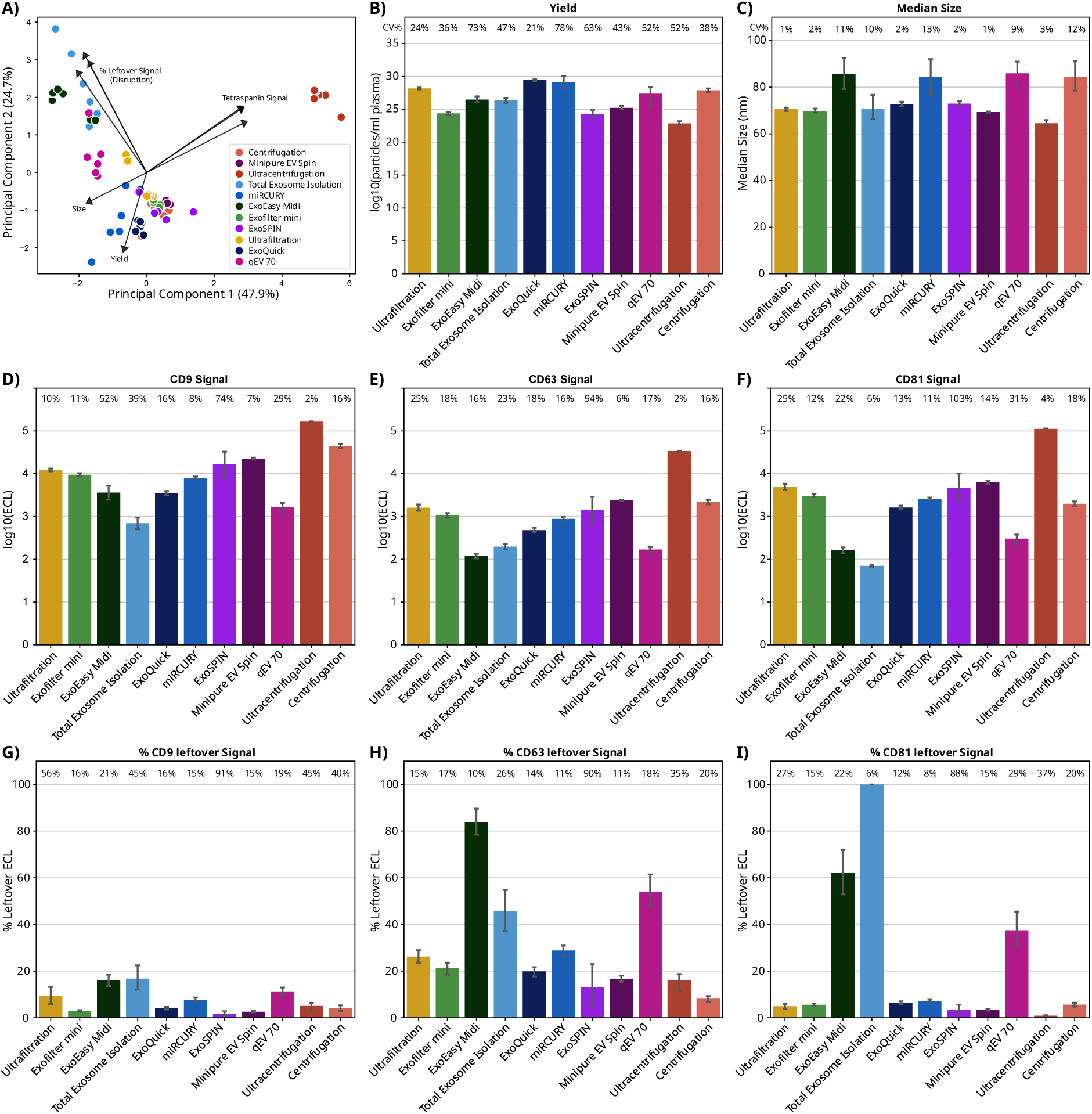
Comprehensive EV characterization: A) Principal Component Analysis (PCA) comparing isolation methods based on particle yield, median size, and tetraspanin signals. B) Particle log10(yields) calculated per mL of input plasma. C) Median diameter (nm) measurements. D-F) Normalized expression of tetraspanins: D) CD9, E) CD63, and F) CD81. Values represent the log10((Electrochemiluminescence (ECL)) normalized to particle count (ECL/1E10 particles). G-I) Percentage of original tetraspanin (CD9, CD63, CD81) signal remaining after detergent treatment (0.25% SDS, 1h). Error bars represent standard deviation from 6 replicate measurements. The coefficient of variation (CV%) for each method is indicated above each bar.

Nano Flow Cytometry analysis showed EV size spanning 65 to 85 nm median (Figure 2C). ExoEasy, miRCURY, and qEV 70 had median diameters of 80-85 nm while Ultra-centrifugation and Centrifugation were smaller at ∼70 nm. Across five donors (Suppl. Figure 3C), centrifugation methods demonstrated the most consistent median sizes, whereas miRCURY, ExoSPIN, and ExoEasy exhibited greater donor-to-donor variability. EV quantification revealed concentrations between 1x10^11^ and 1x10¹³ particles/mL input sample (Suppl. Figure 2A). Particle yield varies by method: precipitation-based methods (miRCURY, ExoQuick) yielded above 1X10¹² particles/mL plasma (Figure 2B), while centrifugation-based approaches yielded approx. 1x10¹^0^ particles/mL. Among donors, yields from centrifugation and miRCURY were more reproducible, while ExoSPIN and ExoEasy showed higher variance (n=5; Suppl. Figure 3B).

Tetraspanin analysis showed distinct patterns across methods in pooled plasma and individual donors (Figure 2D-F, Suppl. Figure 3D-F). CD9 signal/particle ratios varied, with centrifugation methods highest and qEV 70 lowest. CD63 and CD81 followed similar trends; CD63 showed more donor variability, especially in precipitation and affinity-based methods. To confirm tetraspanin signals were vesicle-bound, EV samples underwent SDS detergent treatment, which resulted in significant but method-variable signal reduction (Figure 2G-I, Suppl. Figure 3G-F). Reproducibility varied: Ultra-centrifugation was most consistent (2-4% CV) centrifugation moderate (16-18% CV) and others more variable, particularly Total Exosome Isolation (6-39% CV).

### Proteomics

Proteomics analysis showed differences in protein identification across isolation methods (Figure 3A & Suppl. Table 1). Centrifugation and Ultracentrifugation identified the highest number of proteins (∼1000 each), whereas total Exosome Isolation identified the fewest (161 proteins). The remaining methods identified between 655 and 898 proteins, all surpassing the 487 proteins found in neat plasma. Analysis of individual donor samples revealed minimal inter-donor variation in number of identified proteins within each method (Suppl. Figure 6). Across six replicates, all isolation methods exhibited higher median CV% compared to neat plasma (Figure 3B), with ExoSPIN demonstrating the lowest variability (21%).

**Figure 3:**
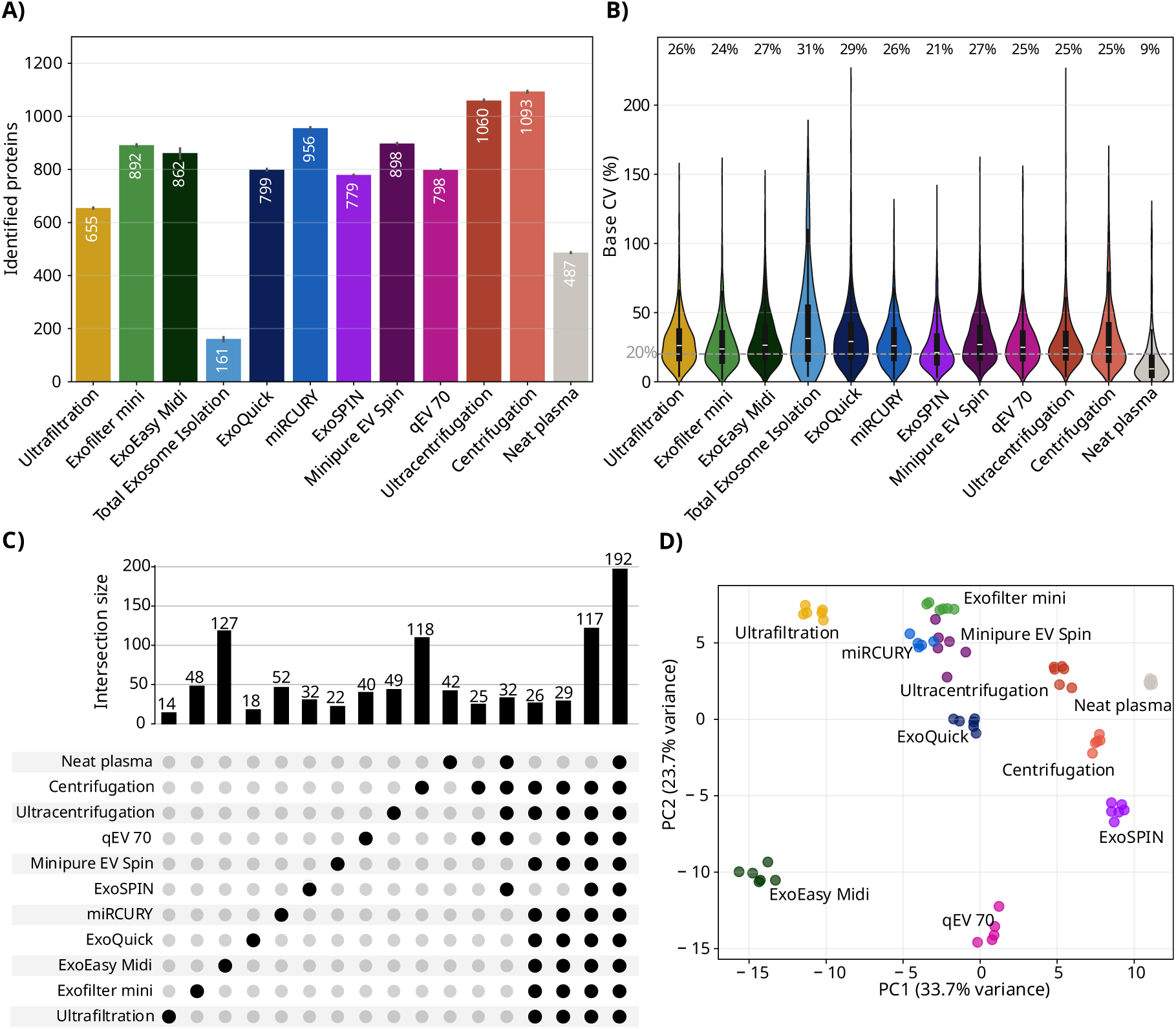
Proteomics analysis of EV isolations. LC-MS/MS based proteomics analysis on pooled plasma, n=6 replicates. (A) Number of identified proteins per isolation method and neat plasma. (B) Distribution of protein-level coefficients of variation (CV) across replicates; violin plots show the full distribution with embedded boxplots. The dashed line marks 20% CV. Annotations report the median CV. (C) UpSet plot illustrating protein intersections across methods. Showing unique protein set for each method, for all other comparisons minimum subset size filtered to n=25 (D) Principal component analysis (PCA) of overlapping proteins (n=192) across methods. Total Exosome Isolation excluded from C and D.

For most methods, more than 50% of the detected proteins did not overlap with neat plasma, and each method identified a unique protein set not observed in other methods (Figure 3C, Suppl. Figure 4 & Suppl. Table 1), with the number of unique proteins ranging from 14 (Ultrafiltration) to 127 (ExoEasy). A core set of 117 proteins was consistently identified across all isolation methods, of these, 69 mapped to the GO term "Extracellular exosome" and included proteins such as heat-shock proteins (HSPA8, HSPA9, HSPD1), histones (H3-7, H3C1, H4C1), and ribosomal proteins (RPL2, RPL12, RPL26).

PCA of overlapping proteins (n=192, excluding Total Exosome Isolation; Figure 3D) separated ExoEasy and qEV 70 from the other methods, while both centrifugation protocols clustered close to neat plasma. PC loadings (Suppl. Figure 7) were dominated by high-abundant plasma proteins, indicating that the PCA mainly reflects plasma carryover rather than vesicle-specific content. PC1 captured mainly immunoglobulin presence, while PC2 captured complement, coagulation and lipoproteins.

All isolation methods enriched low-abundance proteins, while depleting high-abundance plasma proteins compared to neat plasma (Figure 4A, Suppl. Figure 5). Centrifugation-based methods did not reduce albumin (ALB), transferrin (TF), or α1-Acid glycoprotein (ORM1) to the same extent as other methods, with Ultracentrifugation achieving greater depletion than Centrifugation. ExoEasy showed decreased intensities for all the selected proteins relative to neat plasma, which suggests it provides the cleanest depletion of plasma proteins. Ultrafiltration, Exofilter, miRCURY, and Minipure effectively depleted most high-abundance plasma proteins (ALB, ORM1, TF, IGHA1, haptoglobin (HP) and Alpha-2-macroglobulin (A2M) but retained high Alpha-1 antitrypsin (SERPINA1), Fibrinogen alpha chain (FGA) and Apolipoprotein A1 (APOA1), consistent with co-enrichment of lipoproteins and other soluble plasma proteins. Analysis of selected EV marker intensities (Figure 4B) shows that Centrifugation and qEV 70 identified the broadest panel, including CD81, Flotillin 1 (FLOT1), ALIX, CD147 (BSG), HSPA8, HLA A, SDCBP, ITGA2B, and GAPDH [19, 20, 21, 22, 23]. Among tetraspanins also evaluated by MSD only CD81 was detected by LC-MS/MS.

**Figure 4:**
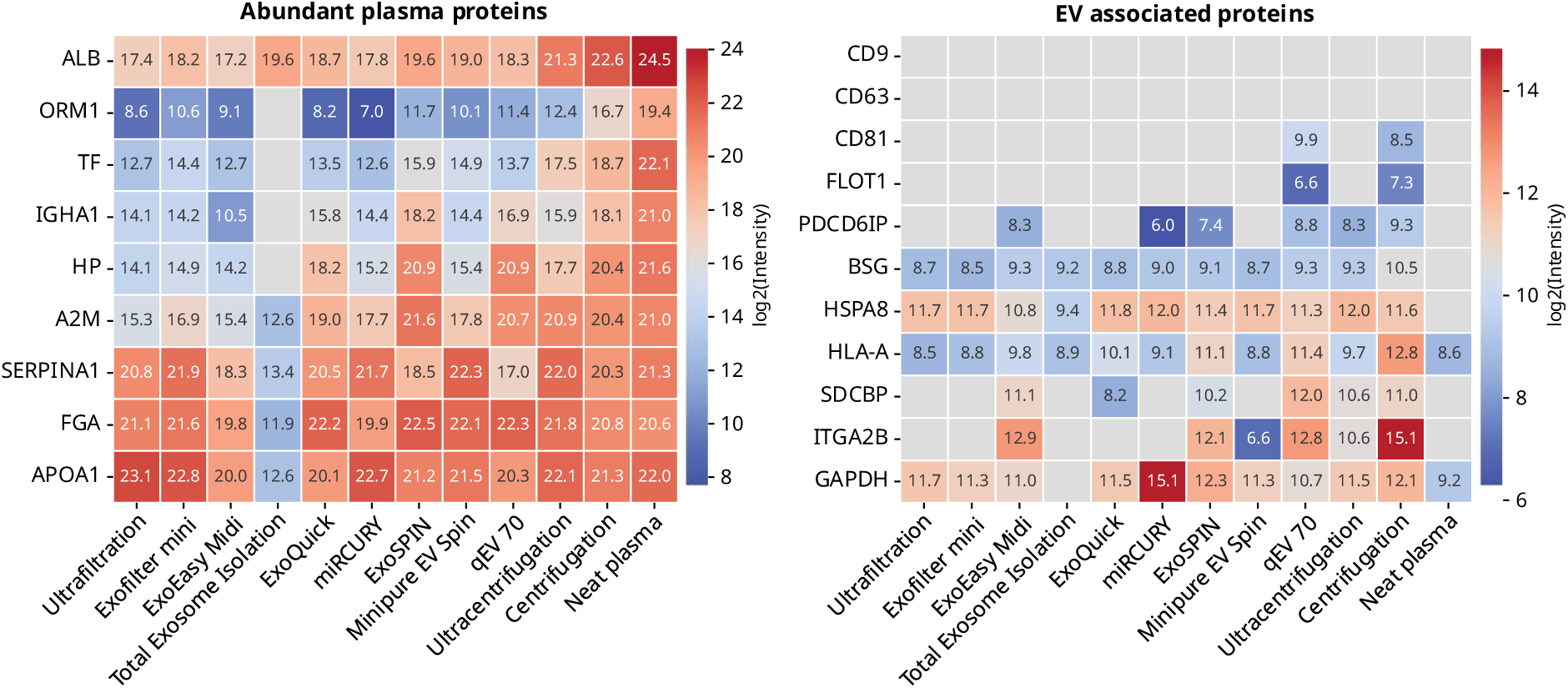
EV and plasma associated protein intensities: (A) Heatmap of log2 transformed intensities of high-abundant plasma proteins for each isolation method and neat plasma (B) Heatmap of log2 transformed intensities of known EV-associated proteins for each isolation method and neat plasma. Pooled plasma, n=6 replicates.

### Evidence-based framework for method selection

To systematically compare the performance of different EV isolation methods, we evaluated each approach across multiple dimensions relevant to downstream applications. Key performance metrics included proteome depth (number of identified proteins by LC-MS/MS), purity (depletion of abundant plasma proteins), particle size distribution, tetraspanin surface marker density, particle yield, reproducibility (coefficient of variation across replicates), and processing time. T These parameters were selected to align with three primary research goals commonly encountered in EV research and relevant to our experimental workflow: achieving high purity for biomarker detection, maximizing proteome coverage for discovery proteomics, or obtaining high particle yields for functional studies. A summary of the comparative performance across all tested methods is presented in Table 2.

**Table 2:** Methods were evaluated using pooled platelet-poor EDTA plasma from 5 healthy donors (n=6 replicates per method). Proteome Depth indicates the number of proteins identified by LC-MS/MS, classified as High (>900 proteins), Moderate (600-900 proteins), or Low (<600 proteins). Purity reflects depletion of high-abundance plasma proteins (ALB, TF, ORM1, IGHA1, HP, A2M, SERPINA1, FGA, APOA1), with ExoEasy Midi showing the most effective depletion (High), while centrifugation-based methods showed minimal depletion (Low). EV Size Range represents median particle diameter measured by NanoFCM, categorized as Small (65-70 nm), Medium (70-80 nm), or Large (80-85 nm) relative to this study. Tetraspanin Signal indicates the density of CD9, CD63, and CD81 on isolated EVs measured by MSD immunoassay (electrochemiluminescence normalized to particle count), classified as High (log1o(ECL) >4.0), Moderate (log1o(ECL) 3.0-4.0), or Low (log1o(ECL) <3.0). CV% values represent coefficient of variation across replicates for tetraspanin measurements, size, and yield, with lower values indicating better reproducibility; ultracentrifugation showed the most consistent tetraspanin measurements (2-4%), while size measurements were highly consistent across all methods (1-13%), and yield variability was highest (21-78%). Particle Yield indicates particles per mL of input plasma: High (>10¹² particles/mL), Medium (10¹¹-10¹² particles/mL), or Low (<10¹¹ particles/mL). Processing Time includes both hands-on and waiting time: Quick (<2h), Moderate (2-4h), or Long (>4h).

| Method | Proteome Depth | Purity | EV Size |  | Tetraspanins |  | Particle Yield |  | Processing Time |
| --- | --- | --- | --- | --- | --- | --- | --- | --- | --- |
|  |  |  | Range (nm) | CV% | Signal | CV% | Yield | CV% |  |
| **Ultrafiltration** | Moderate (815) | Moderate | Medium (70-75) | 1% | Moderate | 10-25% | High | 24% | Moderate (2-3h) |
| **ExoFilter mini** | Moderate (898) | Moderate | Medium (70-75) | 2% | Moderate | 11-18% | Medium | 36% | Quick (2h) |
| **ExoEasy Midi** | Moderate (655) | High | Large (80-85) | 11% | Low | 16-52% | Medium | 73% | Moderate (2-3h) |
| **Total Exosome Isolation** | Low (161) | Low | Small (65-70) | 10% | Low | 6-39% | Medium | 47% | Quick (2h) |
| **ExoQuick** | Low (502) | Low | Medium (70-75) | 2% | Low | 13-18% | High | 21% | Quick (overnight + 1h) |
| **miRCURY** | Moderate (798) | Moderate | Large (80-85) | 13% | Moderate | 8-16% | High | 78% | Quick (overnight + 1h) |
| **ExoSPIN** | Moderate (860) | Moderate | Medium (70-75) | 2% | Moderate | 74-103% | Medium | 63% | Quick (overnight + 1h) |
| **Minipure EV Spin** | Moderate (830) | Moderate | Medium (70-75) | 1% | Moderate | 6-14% | Medium | 43% | Quick (1h) |
| **qEV 70** | Moderate (719) | Moderate | Large (80-85) | 9% | Low | 17-31% | Medium | 52% | Moderate (3-4h) |
| **Ultracentrifugation** | High (1,003) | Low | Small (65) | 3% | High | 2-4% | Low | 52% | Long (6h) |
| **Centrifugation** | High (1,093) | Low | Large (80-85) | 12% | High | 16-18% | Medium | 38% | Moderate (2-3h) |

Our comprehensive characterization provides a decision-guiding framework to assist researchers in selecting the most appropriate EV isolation strategy for their specific research objectives (Figure 5). The framework is organized around three primary research goals: high purity, high proteome coverage, or high particle yield. Each primary goal branches into relevant subcategories that further refine method selection based on specific experimental needs. Not all evaluated methods are represented; only those demonstrating clear performance advantages in decision-relevant categories were included.

**Figure 5:**
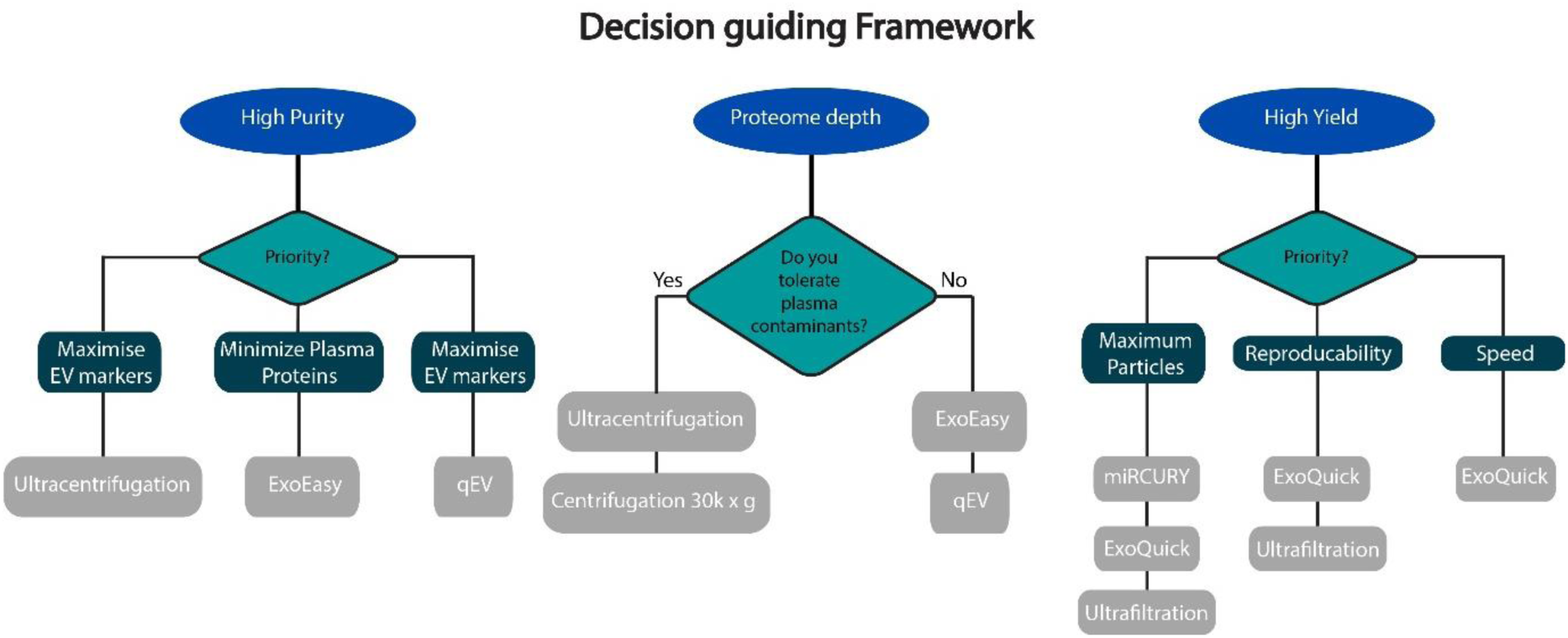
Flowchart guiding method selection based on three primary research goals: high purity, high proteome coverage, or high particle yield. Each pathway branches into subcategories that further refine selection, directing users to the most suitable method(s). Key performance characteristics (proteome depth, purity level, tetraspanin reproducibility, particle yield, and processing time) are listed below each recommended method to guide final decision-making.

## Discussion & Conclusion

We systematically evaluated eleven EV isolation methods across physical properties, membrane markers, and proteomic profiles. Pre-analytical variables and isolation strategies both significantly shaped EV characteristics as well as yield, purity and proteome composition. Isolation methods differed considerably in their ability to retain EV markers, deplete plasma proteins and expand proteome depth. No method excels across all dimensions: Centrifugation methods isolated small EVs with high tetraspanin signals, maximise proteome depth and showed stable reproducibility, but co-isolated plasma contaminants. ExoEasy and qEV 70 isolated larger EVs, with superior plasma protein depletion but lower reproducibility. Ultrafiltration, Exofilter and Minipure captured median-sized EVs, with strong tetraspanin signals, yet retained lipoproteins. Precipitation methods (ExoQuick and miRCURY) yielded high particle counts but poor purity. These distinct method characteristics, emphasise the need to match the isolation approach to the specific scientific objective. We summarized our findings into a flowchart to help guide future studies (Figure 5).

Pre-analytical standardization proved critical. We recommend platelet poor plasma over serum for general EV studies, as serum preparation releases platelet-derived EVs that can overwhelm signals from other cell types [18]. Implementation of an additional centrifugation step effectively depleted platelets, establishing platelet-poor EDTA plasma (PPP) as standard. All samples underwent commonly recommended pre-clearing (0.45 µm filtration and 12,000 × *g* centrifugation) for removal of larger particles and cellular debris. This step enhanced purity but reduced total protein identifications – likely reflecting both contaminant removal and some EV. Omitting pre-clearing achieved proteomic depth comparable to established centrifugation-based protocols [19]. Notably, pre-clearing may be needed when contaminant reduction is critical for downstream assays. Buffer Exchange (Amicon 100kDa MWCO) prior to protein digestion enabled cross-method comparison and minimally impacted pre-cleared samples but reduced protein identification in non-cleared samples. While useful for method comparison, omitting this step is advisable to simplify sample preparation.

EV isolation substantially expanded proteome depth beyond neat plasma, consistent with previous findings [9, 19]. Likely supported by depletion of high-abundance plasma proteins unmasking low-abundant vesicle-associated proteins in a LC-MS/MS workflow. A core set of 117 proteins, including known EV markers, were consistently detected across all methods. In addition, each method captured unique protein subsets, indicating that isolation strategy directly shapes the recovered proteome by selectively enriching specific EV subpopulations and co-isolated plasma components.

Ultracentrifugation, often recognized as the gold standard, efficiently isolated small EVs (median 65nm), consistent with the high g-forces that pellet small, dense particles [16]. MSD profiling effective membrane disruption confirming predominantly vesicular origin with minimal contamination from non-vesicular tetraspanin-containing particles, in accordance with MISEV2018 guidelines [8, 20, 21]. Similarly, Centrifugation (30,000 x *g*,1h) captured small EVs with tetraspanin signals, though at lower recovery than Ultracentrifugation. Yet, proteomics analysis indicated higher plasma protein carryover in centrifugation-based methods compared to the kits we evaluated, likely from co-pelleting lipoproteins or protein/EV aggregation during prolonged spins [3]. Thus centrifugation-based methods present a trade-off, i.e., comprehensive protein coverage with substantial plasma background. There is growing interest in studies of tissue-derived EVs from specific organs, such as liver- and neuron-derived EVs [23, 24]. For such downstream application, including an immunoprecipitation step, it may be less Important with respect to non-EV contaminants as compared to direct analysis of isolated global EVs.

ExoEasy and qEV 70 isolated larger EVs (>80 nm) enhancing overall yield. The disruption assay showed only partial signal reduction, reflecting either greater detergent resistance of larger EVs [22], leftover of cellular components large enough to carry multiple tetraspanins or incomplete removal of non-EV particles/aggregates. Nevertheless, ExoEasy demonstrated superior plasma protein depletion compared to all other methods, coupled with robust EV-associated protein detection. This combination makes ExoEasy particularly advantageous for downstream proteomics analysis.

Recent work by Zhang et al. [9] evaluated nine EV isolation methods from minimal plasma volumes (100 µL), overlapping with four of our methods (UC, qEV, ExoQuick, TEI). Both studies found centrifugation-based methods provided broad proteome coverage with plasma protein carryover and high reproducibility across methods (CV <20%). However, our use of manufacturer-recommended volumes yielded substantially deeper proteome coverage (1093 vs 785 proteins), likely reflecting optimized method performance. Zhang et al. found affinity methods not included my us (MagNet, MagCap) to yield purest EVs, while our ExoEasy showed superior plasma protein depletion among tested methods. Notably, our qEV 70 isolated larger EVs with effective contaminant removal, whereas their qEV 35 showed higher yields but broader size distributions, indicating pore size critically influences captured EV populations Method reproducibility is crucial for reliable EV analysis in clinical applications. Protein intensities showed higher variation (CV 21-31%) after isolation than in neat plasma, reflecting cumulative processing variance. Size measurements showed high consistency across replicates (CV 1–12%), while yields varied more substantially (CV 21–73%), with ultracentrifugation, miRCURY, and Minipure delivering the most stable tetraspanin readouts aligning with prior EV method evaluations [25, 26, 27]. Despite donor-to-donor differences performance patterns remained stable, indicating potential for clinical application when aligned with specific isolation goals. Automation could further improve reproducibility by reduction of manual handling and standardizing processing steps. Our decision framework Figure addresses a fundamental challenge in EV research: no single isolation method optimizes all performance metrics simultaneously. The framework makes these trade-offs transparent: methods maximizing proteome depth compromise purity, while approaches achieving superior contaminant depletion reduce particle recovery. By systematically mapping method performance to experimental priorities, this framework transforms method selection from seeking a universal "best method" into identifying the optimal approach for each specific application, thereby improving reproducibility and facilitating clinical translation of EV research. Study Limitations and Future Directions

The lack of proteomics analysis across different blood collection methods in our analyses leaves our assumption about improved tissue-specific signals through platelet-derived EV depletion unverified. Pre-clearing and buffer exchange, may have limited EV recovery and proteome depth, resulting in lost sensitivity and potentially an incomplete comparative assessment. LC-MS/MS analysis have different analytical sensitivity than other proteomics workflows and that impact the protein IDs we could report in this study. Finally, the absence of clinical metadata limits direct validation for biomarker applications, despite promising reproducibility across donors. Future studies will analyse clinically annotated samples for specific disease investigations, focusing on capturing disease-relevant EV subpopulations and their associated biomarkers.

## Acknowledgement

We thank Jonas Albinus for his valuable contribution to the development of the proteomics workflow. The manuscript was refined and polished with the assistance of the large language models Claude, Gemini and ChatGPT.

