## Supplementary for "No One-Size-Fits-All: An Evidence-Based Framework to Select Plasma EV Isolation Methods"

### No One-Size-Fits-All: Method-Dependent Recovery of Distinct EV Subsets Across 11 Plasma derived EV Isolation Methods

### Supplementary

### Supplementary Figures

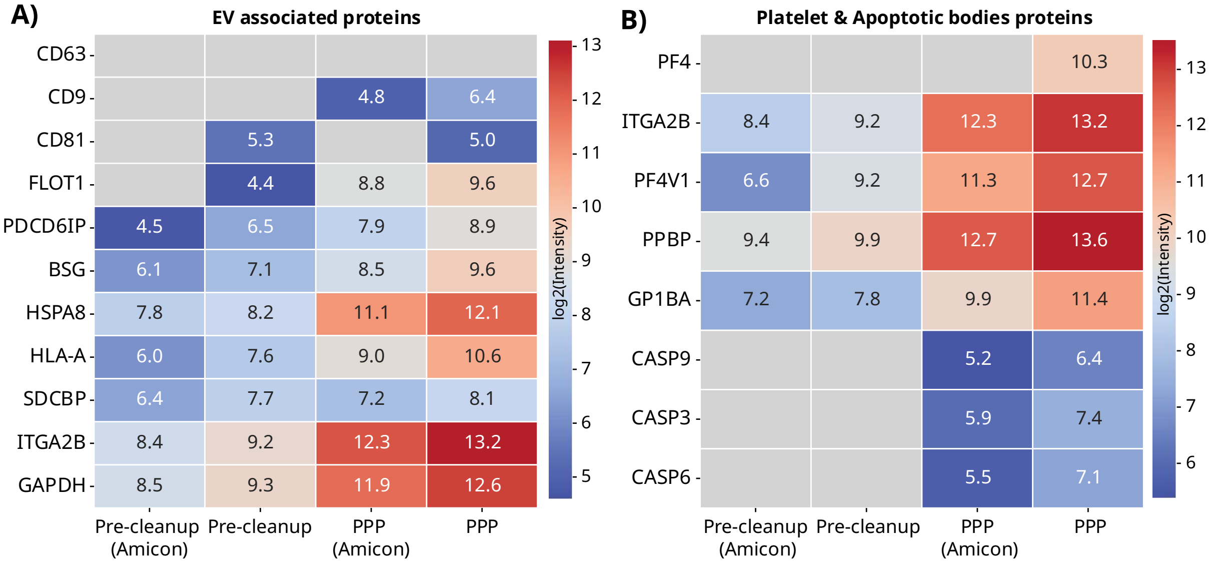

**Supplementary Figure 1: Protein levels after pre-clearing steps:** Heatmaps of LC-MS(MS log2(intensity) of proteins following EV enrichment by centrifugation (30,000 x *g* for 1 hour) from pre-cleared platelet poor plasma (PPP) or non pre-cleared PPP, with and without post-isolation buffer exchange (Amicon) with Amicon Ultra filters (100kDA). (A) EV-associated proteins. (B) Proteins associated with platelets (PF4, ITGA2V, PF4V1, PPBP, GP1BA) and apoptotic bodies (CASP9, CAPS3, CASP6).

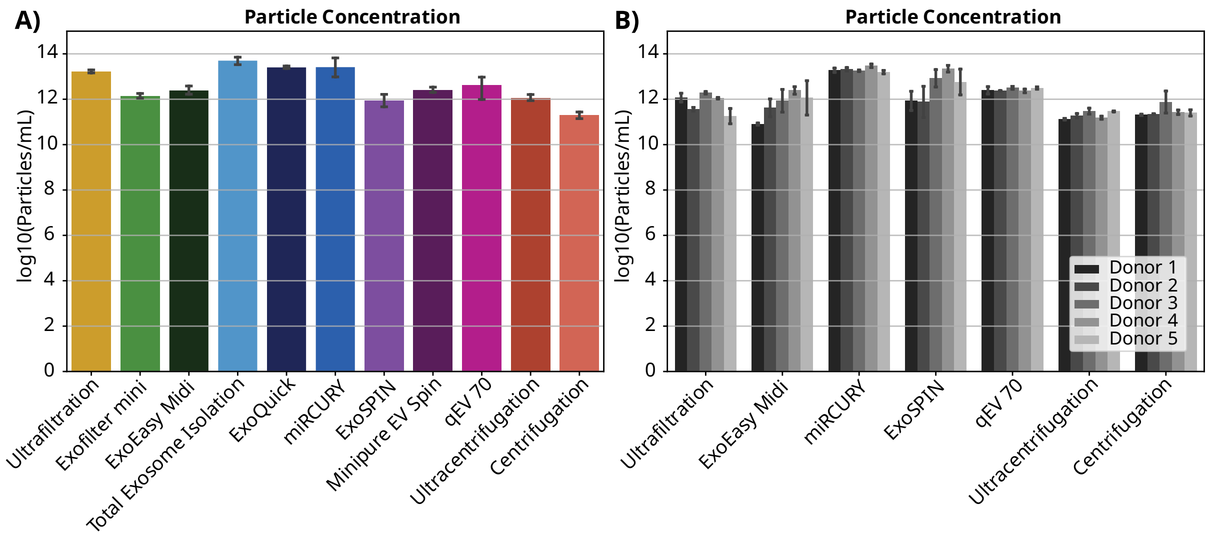

**Supplementary Figure 2: Particle concentration (nanoFCM): (A)** Comparisons of particle concentration across isolation methods in pooled plasma after isolation. **(B)** Particle concentration in samples from selected isolation methods and 5 different donors.

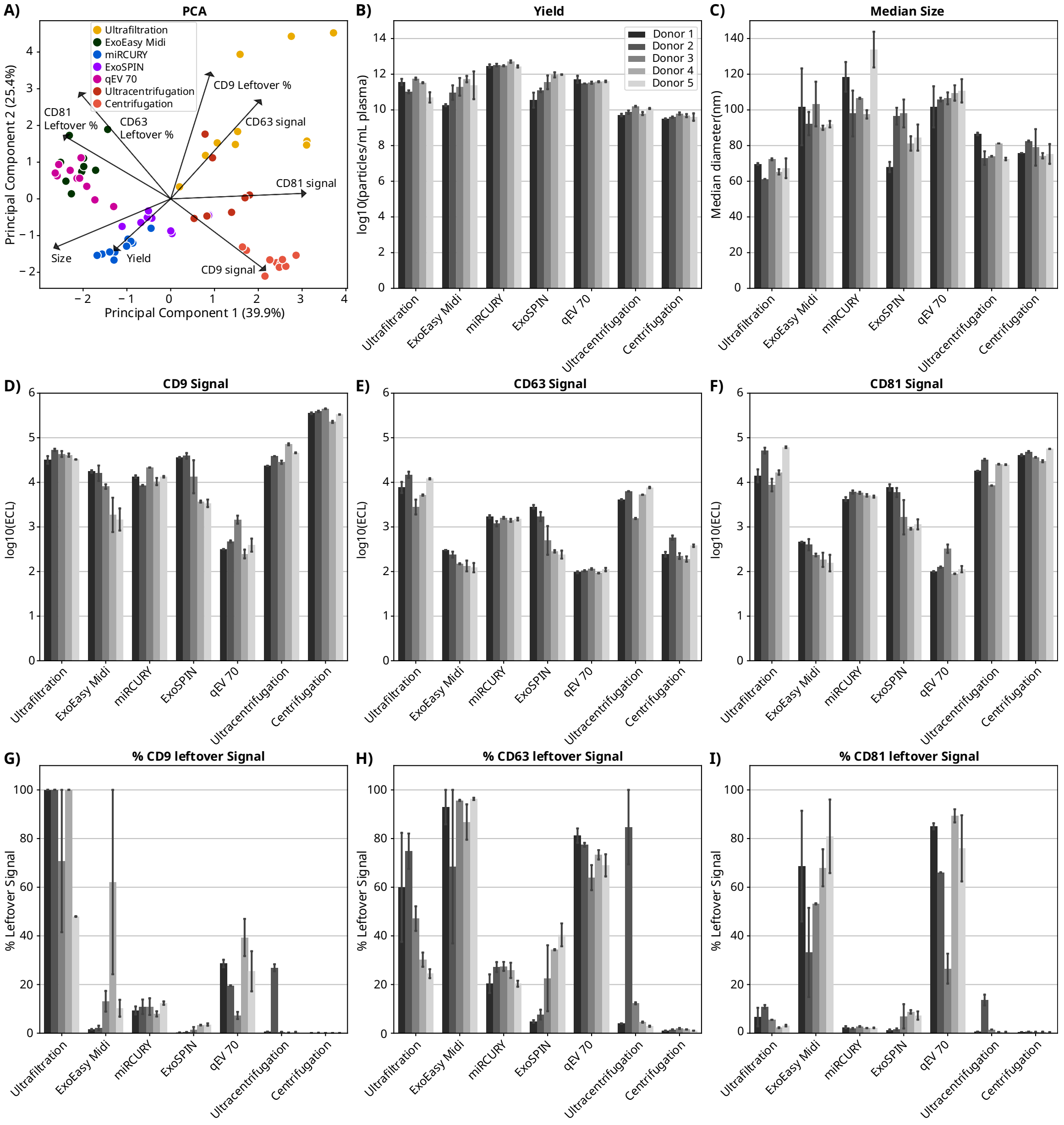

**Supplementary Figure 3: EV characteristics – individual donors:** A) Principal Component Analysis (PCA) of isolation methods based on tetraspanin signals (before and after disruption), particle yield, and median size. B) Particle log10(yields) calculated per mL of input plasma. C) Median diameter (nm) measurements. D-F) Normalized expression of tetraspanins: D) CD9, E) CD63, and F) CD81. Values represent the log10((Electrochemiluminescence (ECL)) normalized to particle count (ECL/1E10 particles). G-I) Percentage of original tetraspanin (CD9, CD63, CD81) signal remaining after detergent treatment (0.25% SDS, 1h). Error bars represent standard deviation from 6 replicate measurements.

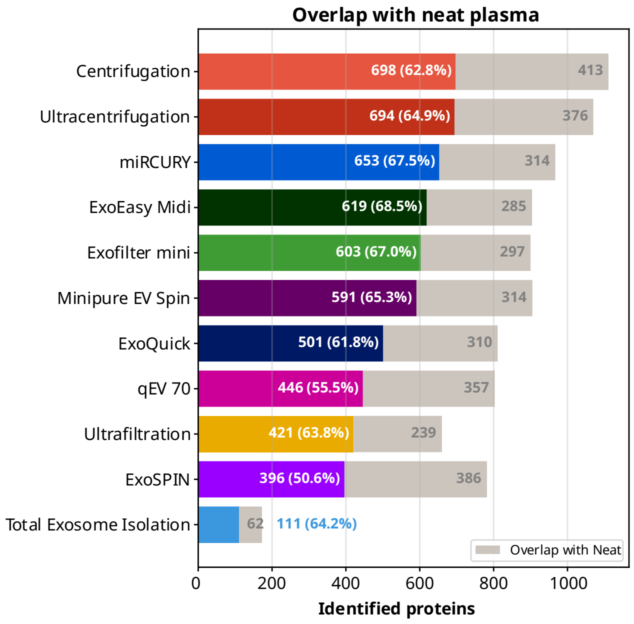

**Supplementary Figure 4: Protein overlap with neat:** Horizontal bars show the total number of proteins identified by each isolation method. Grey bars represent proteins also identified in neat plasma. Coloured bars represent proteins uniquely identified by each method (not detected in neat plasma). Methods are ordered by total protein count.

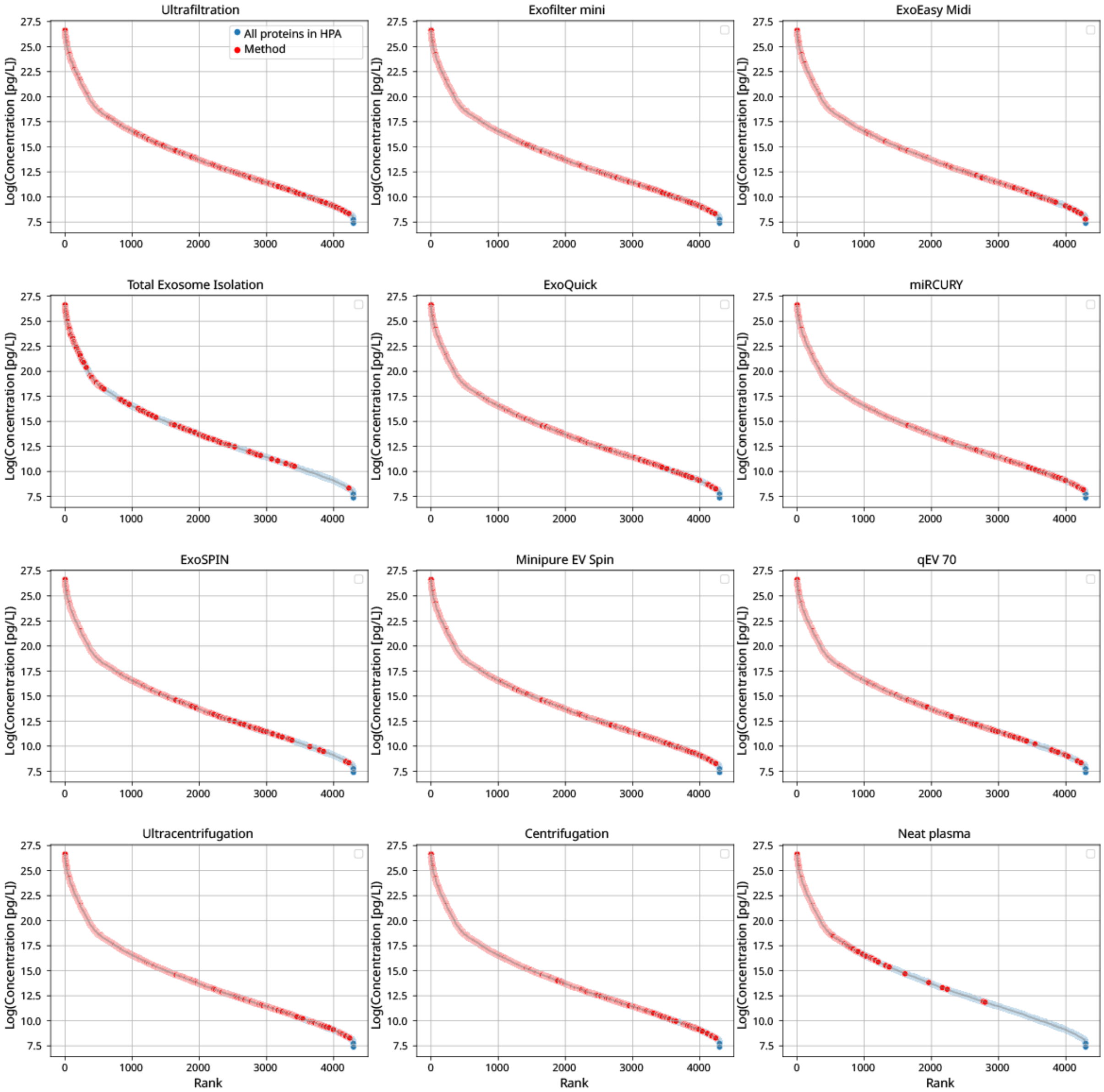

**Supplementary Figure 5: Dynamic range of LC-MS/MS identified proteins:** Distribution of protein concentrations from Human protein Atlas (blue). Proteins are ranked by decreasing concentration (Rank =1 is highest abundance). If a protein is detected by the individual isolation method (pooled plasma n=6 replicates), it is marked (red).

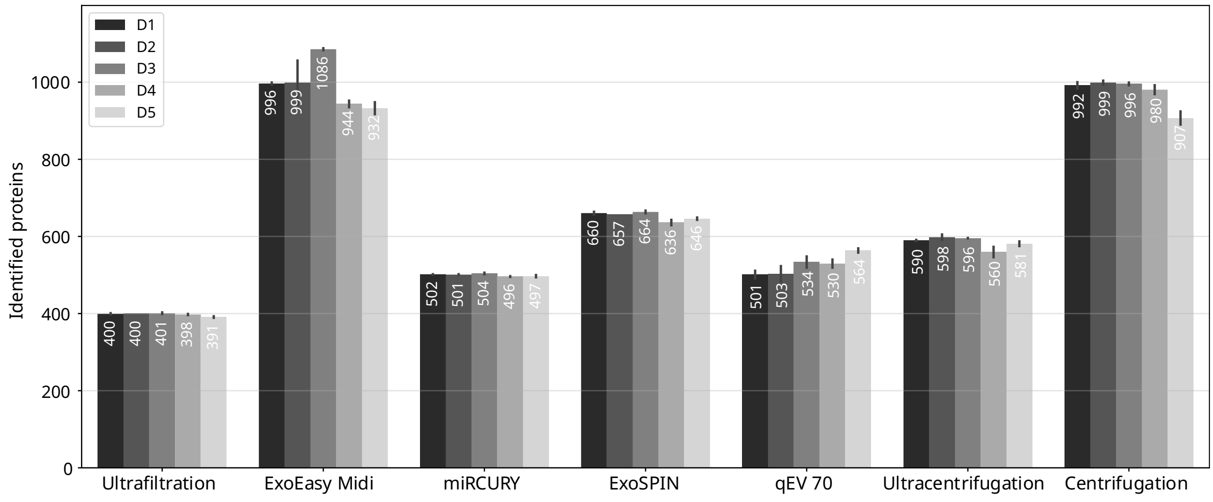

**Supplementary Figure 6: Identified proteins with LC-MS/MS on individual donor samples:** Bars show number of identified proteins for each donor for each isolation method.

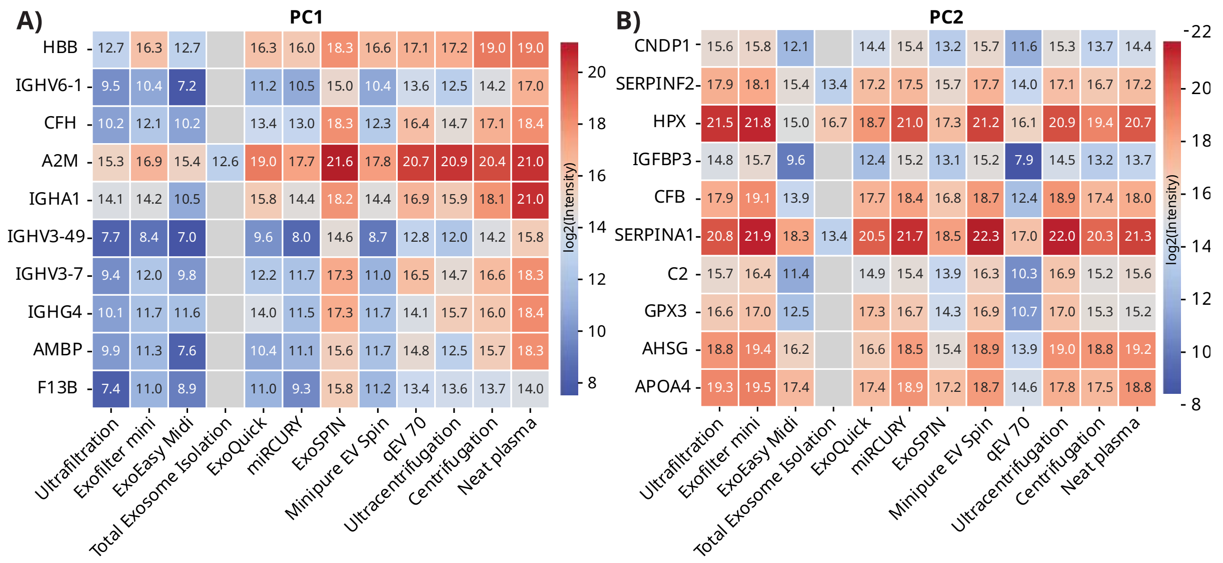

**Supplementary Figure 7: PCA top 10 loadings intensities:** Heatmap of log2 transformed intensities of top 10 PC loadings (proteins), from PCA of overlapping proteins (excluding Total Exosome Isolation; Figure 3D). (A) PC1and (B) PC2.

### Supplementary Tables

**Supplementary Table 1: MSD & nanoFCM results:** Sheet A) MSD results for pre-analytical test MSD signals (ECL/1E10 Particles) for CD9, CD63, CD81, and CD41a, across different blood collection and processing methods (n=5 donors). Sheet B and C) MSD immunoassay and nanoFCM results for each isolation method. MSD signals (ECL/1E10 Particles) for CD9, CD63, and CD81, along with the percent residual signal after disruption relative to no disruption. nanoFCM metrics include median particle size (nm), yield (particles per mL of input plasma), and concentration (particles per mL in the final EV preparation). B) results for EVs isolated from pooled plasma (n=6 replicates), C) results for EVs isolated from 5 individual donor samples (n=3 replicates)

**Supplementary Table 2: Log2-transformed LC-MS/MS intensities (Pooled plasma):** Log2-transformed LC-MS/MS intensities for all proteins identified in isolated EV samples (pooled plasma, n=6 replicates) across all tested isolation methods and neat plasma. The Gene Name column is color-coded by method uniqueness; sorting by colour allows identification of proteins unique to each method.

### Extended Methods

**Ultrafiltration with centrifugation**

Centrifugal filter tubes (Amicon Ultra, 100 kDa MWCO, Cat # UFC510096) were equilibrated with 2,000 µL PBS, then centrifuged at 4,000 x *g* for 10 minutes; flow-through was removed. Samples (500 µL) were diluted 1:5 with PBS, applied to filter units, and centrifuged at 4,000 x *g* for 30 minutes at room temperature. If retentate exceeded 500 µL, further centrifugation was done in 5-minute increments until reaching 500 µL. Samples were washed with 5,000 µL PBS and centrifuged for another 30 minutes under the same conditions. EVs in the retentate were collected and frozen.

**Exofilter mini (HansaBiomed, Cat # 60001)**

Units were activated by pipetting 300µl DI water on the filter and letting it pass through with gravity. After activation, samples (500 µL) were added in a 1:2 dilution with PBS and let flow through by gravity. Subsequently samples were transferred to a tube and eluted using the kit provided elution buffer under gravity. After the sample has passed through, tubes were placed in a tabletop centrifuge (Ole Diech 157) and centrifuged at 500 x *g* for 1 minute for optimized recovery. Eluted samples were transferred to 1.5ml tubes.

**ExoEasy Midi (Qiagen, Cat# 77144)**

Samples (1,000 μL) were mixed 1:1 with XBP buffer, warmed to room temperature, and added to spin columns. They were centrifuged at 500 x *g* for 1 minute. After discarding the flow-through, columns were washed with 4,000 μL XWP buffer and centrifuged at 4,000 x *g* for 5 minutes. Samples were transferred to clean tubes, and 250 μL Elution Buffer XE was added, incubated for 1 minute, and centrifuged at 500 x *g* for 1 minute; the elution was repeated. Eluate was transferred to low-bind tubes.

**Total Exosome Isolation (Thermo Fisher Scientific, Cat# 4484450)**

Plasma was divided into 500 μL aliquots, mixed with 250 μL PBS, and 25 μL Proteinase K, vortexed, and incubated at 37°C for 10 minutes. After adding 155 μL Exosome Precipitation Reagent and incubating at 4°C for 30 minutes, samples were centrifuged at 10,000 x *g* for 5 minutes at room temperature. The supernatant was discarded, and after repeating the centrifugation, pellets were resuspended in 250 μL PBS.

**ExoQuick Ultra (System Biosciences, Cat# EQULTRA-20A-1)**

Plasma (250 μL) aliquots were mixed with 67 μL ExoQuick reagent and incubated for 30 minutes at 4°C. Extracellular vesicles were pelleted by centrifugation at 3,000 x *g* for 10 minutes at room temperature. After removing supernatant, samples were resuspended in 200 μL Buffer B. Purification involved adding 200 μL Buffer A, followed by loading onto prepared kit columns, incubating for 5 minutes at room temperature, and centrifuging for 30 seconds at 1,000 x *g*.

**miRCURY (Qiagen, Cat# 76603)**

Plasma samples (600 μL) were thawed on ice, treated with 6 μL Thrombin (500 U/mL), and incubated for 5 minutes at room temperature. After centrifugation at 10,000 x *g* for 5 minutes, 500 μL of supernatant was mixed with 200 μL Precipitation Buffer A and incubated overnight at 4°C. Samples were centrifuged at 500 x *g* for 5 minutes at 20°C, and pellets were resuspended in 270 μL Resuspension Buffer.

**ExoSPIN (Cell Guidance Systems, Cat# EX02-25)**

Plasma (250 µL) was mixed with Exo-spin™ Buffer in a 1:2 ratio, incubated over night at 4°C, centrifuged at 16,000 x *g* for 30, and the exosome pellet was resuspended in 100 µL PBS. The resuspension was applied to equilibrated Exo-spin™ columns, allowed to flow through by gravity, eluted with 180 µL PBS, and briefly centrifuged.

**qEV 1 70nm (Izon, Cat# ICO-70)**

The qEV1 column with a 70 nm cutoff was used according to the manufacturer's protocol for isolating extracellular vesicles. The column was equilibrated with PBS before loading 500 µL samples. Samples (500 µL) were loaded, and 0.5 mL fractions were collected. Fractions 7–9, enriched with EVs, were pooled. Pooled samples were concentrated using Amicon Ultra filters with a 100 kDa cutoff, topped up with PBS.

**Minipure EV spin (HansaBioMed, Cat# HBM-mPEVS-12)**

Spin columns were equilibrated to room temperature for 30 minutes and then centrifuged at 200 x *g* for 3 minutes to remove the preservative buffer. Samples (200 µL) were then loaded onto the columns, and EVs were eluted by centrifuging at 200 x *g* for 3 minutes.

**Ultracentrifugation**

Isolation was performed by dividing pre-cleared PPP into 200 µL aliquots for ultracentrifugation at 170,000 x g for 2 hours. The pellet was washed in PBS and ultracentrifuged again under the same conditions. The isolated EVs were resuspended in 50 µL of PBS.

**Centrifugation**

Samples of pre-cleared PPP (500 µL) were diluted 1:1 with PBS prior to centrifugation. The mixture was centrifuged at 30,000 x g for 1 hour. The pellet was washed with PBS and centrifuged again under the same conditions.
